# BioCluster: Clustering and Zero-Shot Labeling of Ecological Images Using Foundation Models

**DOI:** 10.64898/2026.10.15.759340

**Authors:** Shakil Ahmed, Kim Bjerge, Toke T. Høye, Christian Nørgaard Storm Pedersen

## Abstract

The increasing automation of biodiversity monitoring is generating large volumes of ecological images requiring taxonomic identification, creating a substantial annotation bottleneck. Assigning species-level labels requires considerable time and expert knowledge, limiting scalability. We propose BioCluster, a framework for unsupervised clustering and zero-shot labeling of ecological images using foundation models. The framework extracts image embeddings from pretrained models and groups images into taxonomic clusters without task-specific training. Species labels are assigned through zero-shot text matching against the taxa expected in the dataset. We investigate dimensionality reduction and six clustering algorithms for organizing fine-grained imagery. A per-image uncertainty score combines clustering and labeling information to sort images into three confidence tiers, so experts can focus on uncertain and unlabeled cases. We evaluate it on two moth and two bird datasets, with one annotated camera-trap collection per taxonomic group. For these camera-trap datasets, per-image species-level accuracy reached 84% on DK-Moth and 83% on HUGO-Bench Aves. An interactive report presents clusters and zero-shot labels by confidence tier and prioritizes cases requiring expert review. The framework provides a training-free approach for organizing and labeling ecological image collections, reducing manual annotation while retaining expert oversight and demonstrating the potential of foundation models for scalable, semi-automated annotation.

## Introduction

Ecological image collections generated by camera traps and other automated monitoring systems can comprise millions of images, creating substantial annotation bottlenecks that limit the scalability of biodiversity research^1–4^. Taxonomic labels must be assigned to these images, either manually by experts or automatically using deep learning classifiers. Both approaches, however, require manually labeled training data, making annotation costly in terms of both time and specialist knowledge. Recent advances in foundation models trained on large-scale visual datasets^5^, as well as models developed specifically for biological applications^6^, provide new opportunities for reducing this dependency on manual annotation. In particular, biological foundation models trained using contrastive learning between images and taxonomic text can associate visual observations with taxonomic concepts and thereby enable zero-shot species recognition without requiring species-specific training data^7^.

BioCLIP^7^ introduced a biological vision-language model trained through contrastive learning between organismal images and hierarchical taxonomic names. By aligning visual and textual representations, BioCLIP enables zero-shot classification of taxa that were not seen during training and can generalize across related taxonomic levels. BioCLIP 2^6^ extended this approach by training on 214 million biological images. The resulting embedding space exhibits emergent biological structure: inter-species variation aligns with functional traits such as beak size and habitat, while within-species variation, including differences in life stage and sex, is preserved in distinct embedding dimensions.

This ability of pretrained visual representations to organize biologically meaningful variation also enables zero-shot clustering, in which unlabeled ecological images are grouped according to species or other biological categories without task-specific training. Lowe et al.^8^ demonstrated that embeddings from self-supervised Vision Transformers can form well-defined clusters for fine-grained biodiversity datasets, including taxa not represented during model training. Such approaches, therefore, provide a means of discovering species-level structures directly from image collections without requiring predefined class labels.

Once clusters have been established, species identities can be assigned through zero-shot text matching. Image embeddings are compared with text embeddings representing candidate species names, allowing the taxonomic label to be assigned to the cluster with the highest semantic similarity without additional model training. BioCLIP^7^, for example, performs zero-shot classification by computing the similarity between image and text representations of taxonomic names.

Recent studies demonstrate the potential of foundation models for both moth and bird monitoring. For moths, BioCLIP 2 substantially outperformed other foundation models on Danish moth species from the AMI camera system, while knowledge distillation from BioCLIP 2 into a lightweight ConvNeXt model achieved comparable accuracy at a substantially lower computational cost^9^. For birds, BirdSAT^10^ demonstrated that self-supervised Vision Transformers enhanced with metadata can achieve state-of-the-art fine-grained classification on the bird subset of iNaturalist-2021. TaxaBind further demonstrated strong zero-shot bird classification on Birds525 and CUB-200-2011, outperforming BioCLIP by distilling multimodal information into an image encoder^11^.

In this paper, we develop a framework to cluster ecological images with zero-shot labeling. The framework is evaluated end-to-end on four datasets spanning moths and birds, consisting of curated collections and camera-trap imagery. Our main contributions are as follows:

- A pipeline that combines foundation models, unsupervised clustering, and zero-shot text matching to transform an unlabeled image collection into groups assigned species-level labels using a list of candidate species names.
- A configurable uncertainty score that integrates four complementary signals for each image, along with two configurable thresholds for routing images to automatic acceptance, expert review (an expert checks the proposed label), or human required (an expert identifies the species, because the image has no reliable label).
- The DK-Moth dataset comprises 248 Danish moth species recorded using automated light traps over two summer seasons and curated to the species level by a domain expert.

Together, these developments suggest that combining pretrained visual embeddings for unsupervised clustering with vision-language matching for zero-shot labeling can provide a training-free approach to taxonomic annotation. Such a framework presented in this paper is particularly attractive for biodiversity monitoring, where large image collections, high species diversity, and limited access to expert annotators make conventional supervised classification costly to develop. Human-in-the-loop tools, such as ours, are also well suited to filling taxonomic gaps in training data.

## Methods

### Framework Architecture

Our proposed framework takes a series of steps shown in Figure 1. The initial step in this process is to pass each image through a vision-language foundation model to extract the feature embedding vector that contains both visual and semantic data about the organism. Second, these high-embedding vectors are passed through dimensional reduction methods and then clustered using different clustering algorithms, where images of the same species naturally cluster in the embedding space, and the clustering quality is measured. Third, clusters are then assigned a species-level name by zero-shot text matching, with the cluster centroid compared to text embeddings of candidate species names augmented with taxonomic context. Fourth, a combined uncertainty score is determined based on four independent signals, the outcome of which is then used to mark all annotations at three tiers of confidence. Lastly, a web interface allows a domain expert to verify, correct, or reassign annotations, and these human-in-the-loop decisions are merged back into the final dataset with complete provenance.

**Figure 1.**
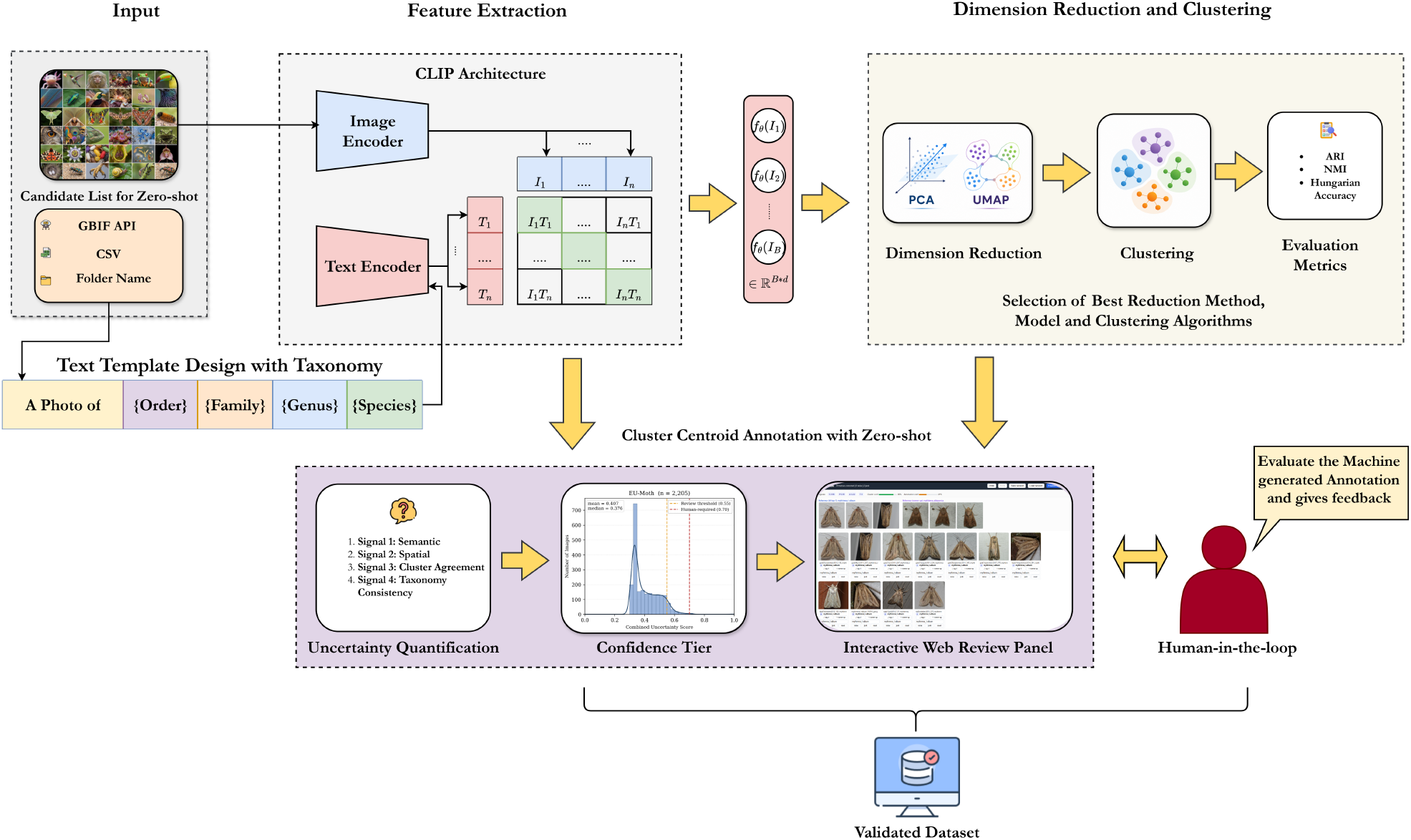
The methodology diagram for clustering and annotation. The framework takes images as input and passes them through the model-specific preprocessing stage. Then it extracts the feature vectors according to the model’s embedding space and passes through the dimensionality reduction phase. The reduced features are then fed into the clustering algorithms. Then, the best chosen model and clustering algorithm perform the zero-shot annotation of the clusters and per species.

### Image Embeddings

We used BioCLIP 2^6^, a domain-specific vision foundation model, as our primary encoder. It is trained using hierarchical contrastive learning on the TreeOfLife-200M dataset and spans more than 950,000 taxa, paired with structured taxonomic text. In particular, we used the largest ViT-H/14^12^ backbone, which was initialized from a LAION-2B CLIP checkpoint because of its recent release^6^. We used the model in inference mode, so no weights were modified, and no task-specific training was performed. Each image passed once through the model’s native preprocessing transform (224 *×* 224 input, identical to the transform used in pretraining) and produced a 1024D embedding.

We write *f*_*θ*_ for the image encoder and *g*_*φ*_ for its text encoder. Images are processed in batches. For a batch of images *{I*_1_, *I*_2_, …, *I*_*B*_*}*, the forward pass through *f*_*θ*_ produces a matrix of raw embeddings,

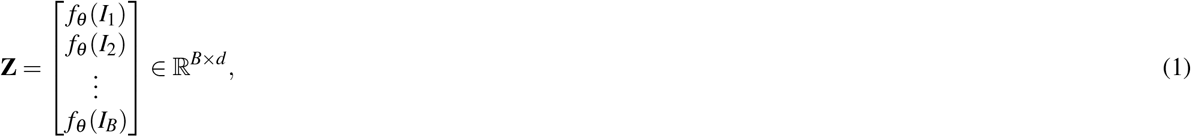

where *d* is the embedding dimension of the model (*d* = 1024 for the primary BioCLIP 2.5 model). Each raw embedding **z**_*i*_ is then L2-normalized onto the unit hypersphere,

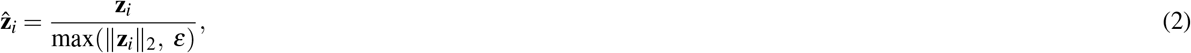

where *ε* = 10^*−*8^ avoids division by zero in the degenerate case of a zero vector. After normalization, every embedding lies on the unit hypersphere in ℝ^1024^, and the cosine similarity between two embeddings reduces to their dot product. This normalization is required by the text-matching stage, which scores matches by cosine similarity.

We also evaluated five further vision foundation models under an identical extraction interface to see how the pipeline behaves across backbones of different scales, domains, and training objectives. The first was the early release of BioCLIP 2 (768D)^6^, which isolates the effect of backbone scale within the same model. The others were the earlier BioCLIP (512D)^7^, the general-purpose CLIP (768D)^13^, the self supervised DINOv2 (1024D)^5^, and the DINOv3 (1280D)^14^. Each encoder was loaded with its own preprocessing transform, so every model received inputs matched to its own training distribution. DINOv2 and v3 have no text encoder, so those models only participated in the clustering comparison and not in the annotation.

### Dimensionality Reduction Before Clustering

Foundation-model embeddings are high-dimensional. In this regime, distance concentration erodes the contrast that distance-based and density-based clustering relies on^15–17^. We therefore applied a single reduction step before clustering, shared across all clustering methods. The reduction method is configurable and has four modes. The *none* mode keeps the full embedding vector. PCA^18^ keeps the directions of greatest variance and discards low-variance noise, which sharpens the geometry that distance-based and variance-based methods use. UMAP^19^ builds a weighted nearest-neighbor graph and preserves its topology in the reduced space, which helps when the discriminative structure is non-linear. The two-stage mode runs PCA to an intermediate dimension and then UMAP, so the linear step removes noise and speeds up the graph construction before the non-linear step.

Before clustering, we reduced the dimension of each embedding. The embeddings of the six backbones have between 512 and 1,280 dimensions. Allaoui et al.^20^ applied UMAP as a preprocessing step, which improved k-means, agglomerative clustering, GMM, and HDBSCAN by enhancing accuracy and reducing runtime. It also enhanced cluster detection by DBSCAN while reducing its sensitivity to parameter choices^21^. Lowe et al.^8^ found that embeddings from pretrained image encoders were typically clustered best after a manifold-based reduction, and that with UMAP the number of target dimensions mattered little between 5 and 200; they applied this reduction when clustering unseen datasets. On camera-trap images, Markoff et al.^22^ found that reducing the embeddings to two dimensions with t-SNE raised the V-measure of HDBSCAN from 0.498 to 0.783, averaged over five backbones. We used UMAP alone and PCA followed by UMAP^19^, which has no computational restrictions on the output dimension and is therefore usable as a general-purpose dimension reduction for machine learning, whereas t-SNE was introduced to give each point a location in a two- or three-dimensional map for visualization^23^. UMAP has also clustered better than PCA as a preceding reduction, both on document embeddings^24^ and on camera-trap images^22^.

The target dimension also matters. Markoff et al.^22^ clustered the embeddings with two dimensions and noted that this discards substantial information from the original embeddings and may not be optimal for clustering; as future work, they suggested clustering after a higher-dimensional reduction. Our pipeline follows this separation. Clustering is performed in 5 to 20 dimensions, within the range in which Lowe et al.^8^ found the number of UMAP dimensions to matter little, and a separate two-dimensional UMAP projection is computed only for visual inspection. The dimension used for each dataset is given beneath its name in **Supplementary Material Table S1**.

The reduction was applied only to the clustering algorithms and never to text matching. Because zero-shot annotation compares each image with species-text embeddings using cosine similarity, which requires both modalities to share the model’s joint image–text space^6,13^. Therefore, projecting out of that space may destroy the learned alignment.

### Clustering

Clustering groups images of the same species without supervision. We used six clustering algorithms from five different paradigms to find which one best recovers species structure from the embeddings. In our experiment, we considered K-means^25,26^, Spectral^27^, Agglomerative^28^, and Gaussian Mixture Model (GMM)^29^ as fixed-K methods. These methods assign each image to a single cluster, known a priori as the number of clusters *K*, without noise labels. In our experiment, we also considered density-based clustering algorithms such as DBSCAN^30^ and HDBSCAN^31,32^. These methods discover cluster boundaries, learn cluster size *K* using the data, and explicitly indicate low-density points as noise. These findings are reported in the Results section.

For the annotation pipeline, however, we used a single density-based method, HDBSCAN. It generalizes DBSCAN by replacing the single global density threshold (*ε*) with a hierarchy built across all density scales and then extracting the clusters that persist over the widest range of densities.

Three properties make HDBSCAN the preferred clustering algorithm for label-free annotation, and together they explain why we prefer it over fixed-*K* methods. The first is the per-point membership probability *p*_*i*_ *∈* [0, 1], which measures how firmly a point belongs to its cluster. This probability is used directly as the spatial uncertainty signal below, and no fixed-*K* method produces an equivalent. The second is explicit noise labeling. Low-density points are marked as noise rather than being forced into the nearest cluster. In a biological collection, these points are genuinely ambiguous, under-sampled, or low-quality specimens, and routing them to expert review is more defensible than assigning them automatically. The third is density adaptivity. Because HDBSCAN works at local density scales, it recovers both tight and diffuse species clusters in the same dataset without per-cluster tuning.

HDBSCAN infers the number of clusters from the data. We set `min_cluster_size` by a per-dataset sweep that maximizes a coverage-weighted score, ARI *×* (1*−* noise fraction), where ARI is the Adjusted Rand Index^33^ defined under Evaluation Metrics below. This score penalizes degenerate solutions that reach high agreement by discarding most images as noise. The `min_samples` parameter controls how conservative the density estimate is. It is configurable: we take `min_cluster_size` as its starting value, but we adjust it per dataset when the data require it. The cluster-selection method, excess of mass or leaf, is likewise chosen per dataset. Noise points are excluded from the clustering-quality metrics so that an algorithm is not penalized for correctly abstaining. However, they are still annotated and shown in the review interface.

### Zero-shot Annotation

After clustering the species, each cluster is assigned a species name through zero-shot text matching in the original dimensional space of the model, using the vision-language alignment learned during contrastive pretraining^6,7^. We first assembled a candidate species list from one of three interchangeable sources: a regional query to the Global Biodiversity Information Facility (GBIF) occurrence API^34^, a user-supplied CSV file listing the candidate species with their genus, family and order, or the sub-folder names of a curated dataset.

Each candidate species is represented by a single, taxonomically structured prompt of the form `“a photo of {order} {family} {genus} {name}”`, filled from the taxonomy of that species and encoded by the text encoder *g*_*φ*_. The resulting vector is L2-normalized to give the species text embedding **ê**_*s*_. Ordering the ranks from coarse to fine matches the structured taxonomic text on which the BioCLIP family was trained. We also tried an ensemble of several templates whose normalized embeddings are averaged, as is common in CLIP zero-shot classification^13^, but on our data, the ensemble showed no measurable gain over the single structured prompt; indeed, there was a small decrease in accuracy, so we used the simpler single-prompt form. When a taxonomic rank is unavailable for a species, it is omitted from the prompt.

A cluster is annotated by comparing its centroid *µ*_*k*_, the mean of its member embeddings 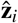, against every species text embedding **ê**_*s*_ using cosine similarity:

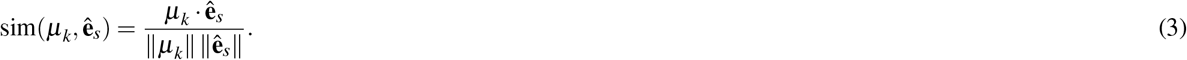

The top-ranked species,

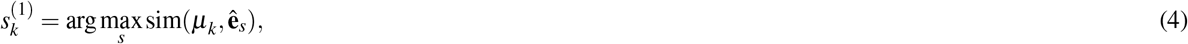

is assigned to all images in the cluster. We also keep the label with the second highest support (runner-up)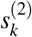, and the margin between the two, 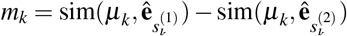, as a confidence measure. In parallel, every image casts an independent vote 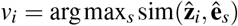. These per-image votes are not used as annotations because the centroid assignment is more stable, but they feed into the cluster-agreement signal below.

### Uncertainty Quantification

Every image receives a composite uncertainty score that combines four independent signals, each aimed at a distinct failure mode. All four signals lie in [0, 1], and higher values indicate lower reliability. A correct annotation is usually low on all four, while different error types raise different subsets.

*Signal 1. Semantic Uncertainty (text margin):* This signal measures how separable the top species is from the runner-up in the text-matching space:

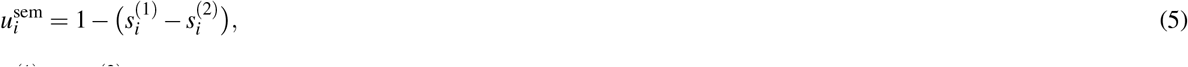

where 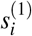 and 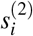 are the two highest cosine similarities of image *i* to the candidate species embeddings. It flags confusion between visually similar species, where the absolute similarities are high, but their differences are small.

*Signal 2. Spatial Uncertainty (membership):* This signal uses the HDBSCAN membership probability *p*_*i*_ introduced above, 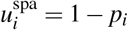, and noise points are given the maximum value of 1.0. If a clustering method other than HDBSCAN is used, the per-sample silhouette coefficient sil_*i*_ *∈* [*−*1, 1]^35^ serves as a proxy, rescaled to the unit interval as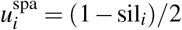.

*Signal 3. Cluster Agreement:* This signal measures the internal voting consistency of a cluster. Let *f*_*k*_ be the fraction of images in cluster *C*_*k*_ whose votes match the cluster’s majority species. The base value is 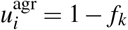. Images that dissent from 2, and noise points receive 1 the majority receive an additional penalty *λ*, 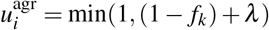, where *λ* = 0.2, and noise points receive 1.0. Without this penalty, the value would be constant across a cluster and would repeat information already provided by the clustering. The increment was chosen heuristically rather than optimized, as larger values primarily trade off auto-accept accuracy against review workload, and it is configurable.

*Signal 4. Taxonomic Consistency:* This signal penalizes clusters whose per-image votes span more than one taxonomic family:

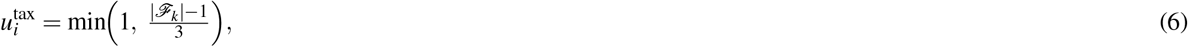

where *F*_*k*_ is the set of distinct families voted within cluster *C*_*k*_. A taxonomically pure cluster receives zero, and the divisor determines where the signal saturates, so a cluster spanning two families scores 1*/*3, three families 2*/*3, and four or more reaches the maximum of 1. The signal needs family-level taxonomy, and it returns zero when that information is unavailable. This was the case for CUB-200-2011 and HUGO-Bench Aves, whose common-name labels do not resolve to a family; therefore, on those two datasets, the combined score rests on the remaining three signals. Noises still receive the maximum value on this signal, so the floor that keeps them out of the auto-accept tier is unaffected.

The four signals are then combined using a weighted average.

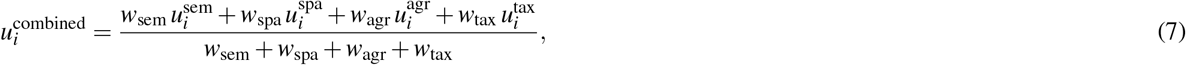

with weights *w*_sem_ = 0.35, *w*_spa_ = *w*_agr_ = 0.25, and *w*_tax_ = 0.15. The pipeline is unsupervised, so no labeled validation set is available to learn these weights. We therefore set them with the Simple Multi-Attribute Rating Technique^36^. The semantic margin receives the largest weight because it is the only signal defined directly in the space where the annotation decision is made. The spatial and agreement signals measure complementary aspects of cluster quality, and neither is clearly more informative than the other, so they are weighted equally. The taxonomic signal receives the smallest weight since it depends on external data that may be absent. Although all weights are configurable.

The combined score is then mapped to three confidence tiers using two absolute thresholds:

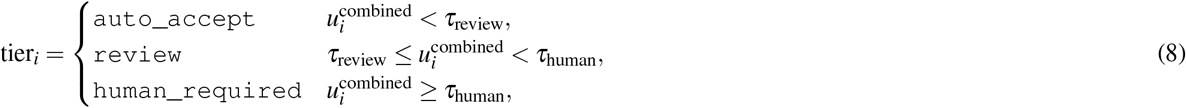

with defaults *τ*_review_ = 0.55 and *τ*_human_ = 0.70. This two-threshold evaluation extends the classical single-threshold reject option^37,38^. We used absolute thresholds rather than percentile thresholds on purpose. A percentile threshold always flags a fixed fraction of images, regardless of quality. An absolute threshold allows a strong clustering to raise few flags, while a weak clustering raises many, which is a more informative measure of reliability. Moreover, these thresholds are set so that an image reaches the review tier only when several signals are elevated together, rather than on the strength of any one of them. They can be lowered when more review capacity is available, which sends more images to an expert, so adjusting them is a trade-off.

Therefore, all of these values are configurable. They do not change the species labels, which follow from the cluster-to-text matching, but they do determine which images reach each tier and hence the accuracy observed within tiers.

### Human-in-the-Loop

Species names are assigned to the cluster level rather than to individual images. Thus, when an expert confirms or corrects a cluster assignment, the species label is resolved for all images within the cluster through a single decision. Only images flagged by the confidence tiers require further expert attention, while individual images remain available for inspection when needed. A self-contained HTML report shown in Figure 2 presents each cluster as a card. The card shows the member images, reference images for the predicted species and the runner-up, the four signal breakdowns, and two separate confidence measures. The clustering-coherence measure,

**Figure 2.**
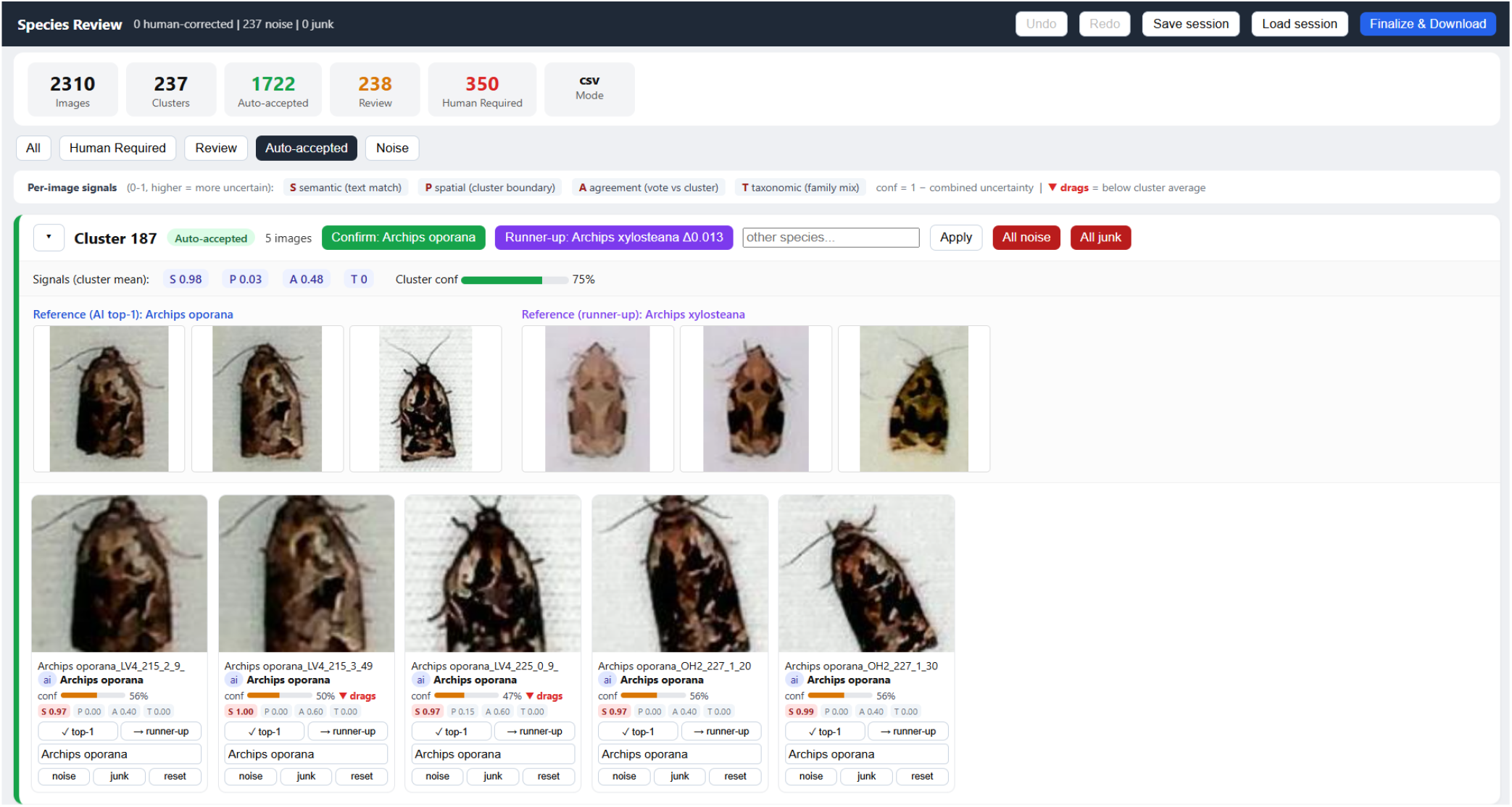
Self-contained interactive HTML report. The HTML report contains photo cards of every images. The report facilitates full flexibility for the human expert. The cluster confidence meter tells how tight the cluster is and the per image annotation meter says how confidence the model to predict the species.

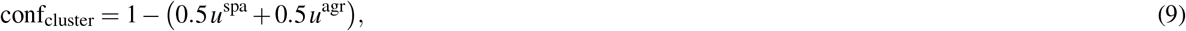

asks whether the images belong together. On the other hand, the annotation-correctness (per-image) measure,

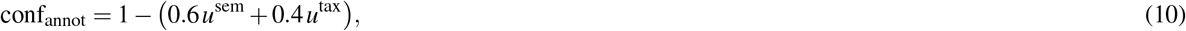

asks whether the assigned name is correct. The clustering meter weights its two signals equally, as both directly measure cluster quality. In contrast, the annotation meter prioritizes the semantic signal, which directly assesses naming correctness, while the taxonomic signal serves as an indirect check. When no family can be resolved, the semantic signal alone determines this meter, since allocating a fixed portion of the scale to a signal that is identically zero would be inappropriate. Separating the two meters enables a human expert to distinguish between a coherent but misnamed cluster and an incoherent one. The expert can then confirm, reassign, or reject at either the cluster or image level, with full redo and undo functionality. Thus, this functionality is specific to the web report interface and does not affect the results.

Human expert decisions are then merged into the final labels, and each label carries a provenance tag. These tags keep the automated and human contributions auditable. The output is organized as a species-labeled image directory, which can be used directly for downstream supervised training or as input to a further iteration of the pipeline.

The web report is a single offline file in which all decisions are made in the browser; because it embeds a thumbnail of every image (transferable HTML file), its size grows with the collection (26.7 MB for one 2,310-image DK-Moth subsample). Therefore, very large collections are better reviewed in the desktop application, which loads images on demand.

### Datasets

We evaluated the pipeline on four benchmark datasets presented in Table 1, chosen to vary the axes that matter for real-world biodiversity monitoring: taxonomic domain, class balance, scale, and image quality. Two of them cover moths (Lepidoptera) and two cover birds (Aves).

**Table 1.** Summary of the benchmark datasets. The two subsampled datasets, DK-Moth (sub) and HUGO-Aves (sub), are each drawn under ten random seeds. Their image counts are reported per run, and all results are averaged over the ten runs. The full HUGO-Bench Aves dataset from which the subsample is drawn holds 73,528 images of the same 30 species.

| Property | EU-Moth | DK-Moth (full) | DK-Moth (sub) | CUB-200-2011 | HUGO-Aves (sub) |
| --- | --- | --- | --- | --- | --- |
| Taxonomic order | Lepidoptera | Lepidoptera | Lepidoptera | Aves | Aves |
| Classes | 200 | 351 | 248 | 200 | 30 |
| Total images | 2,205 | 18,999 | 2,310 | 11,788 | 6,000 |
| Min / class | 7 | 1 | 5 | 41 | 200 |
| Max / class | 14 | 1,729 | 10 | 62 | 200 |
| Mean / class | 11.0 | 54.1 | 9.3 | 59.0 | 200 |
| Class balance | Balanced | Long-tailed | Balanced | Balanced | Balanced |
| Image source | Citizen science | Camera trap | Camera trap | Flickr | Camera trap |
| Naming | Scientific | Scientific | Scientific | Common | Common |
| Runs (seeds) | 1 | 1 | 10 | 1 | 10 |

EU-Moth^39^ contains 2,205 citizen-science images of 200 Central European moth species. It is approximately balanced, with 7 to 14 images per class and uses scientific binomial labels.

DK-Moth contains 18,999 automated camera-trap images of 351 Danish moth species, with a strongly long-tailed distribution: the largest class has 1,729 images, 32.8% of classes have five or fewer images, and the median class size is 15. Its lighting, pose, and image quality are far more variable than in the curated sets. It is the most realistic and, for clustering, the most difficult of the four datasets. DK-Moth has not been published before. We release the balanced 248-species subsamples with this work, the ten draws used in our experiments, to support reproducibility. The full long-tailed collection is described here for context but is not part of this release. The images were collected with an automated light-trap camera system during the 2022 and 2023 summer seasons at three Danish nature areas, Lille Vildmose, Ovstrup Hede, and Søholt Storskov, following the acquisition procedure of Bjerge et al.^4^. The cameras recorded at ten-minute intervals, and individual moth crops were extracted from the full trap images and then manually sorted and curated to the species level by a domain expert (Flemming Helsing-Nielsen).

The other two datasets are birds, and together they probe cross-domain generalization from moths to birds. CUB-200-2011^40^ contains 11,788 Flickr images of 200 North American bird species, balanced at 41 to 62 images per class. Its labels are English common names rather than scientific binomials, so it tests whether text matching tolerates vernacular naming.

The Aves partition of HUGO-Bench^22,41^ is a camera-trap dataset built for zero-shot species clustering. It contains 73,528 images of 30 bird species, also labeled with English common names, and its full distribution is imbalanced, ranging from 1,240 to 4,420 images per class. Pairing it with CUB-200-2011 holds the taxonomic domain fixed while the acquisition conditions change, transitioning a curated dataset to camera trap photographs. It also comes from a published clustering benchmark on the same 30 species, which allows us to compare our clustering quality against an external reference.

Two of the datasets are evaluated through balanced subsamples rather than in full. Each subsample is drawn under ten different random seeds, so that every result on these datasets is reported as a mean over ten runs, and its run-to-run variability can be quantified. For the DK-Moth dataset, the subsampling separates the effect of class imbalance from that of the camera-trap imaging conditions. We kept all eligible 248 classes that have at least 5 images and drew up to 10 images from each, which gives 2,310 images per run. A class with more than 10 images contributes exactly 10 images, while a smaller class contributes all it has, so the class set and the image count are identical across seeds. Only the images that are drawn change for the 192 classes that have more than 10 images. Each seed is thus a repeated draw over the same 248-species. For HUGO-Bench Aves, whose full partition is too large to cluster repeatedly, we sampled exactly 200 images from each of the 30 classes, maintaining 6,000 images per run. The 200-per-class draw also matches the evaluation protocol of the original study by Markoff et al.^22^, which keeps the external comparison on equal terms.

## Evaluation Metrics

We evaluated how well the pipeline clustered images and how reliably it annotated those clusters. The clustering quality is reported with the Adjusted Rand Index (ARI)^33^, Normalized Mutual Information (NMI)^42^, and Hungarian accuracy^43^. When ground-truth labels were available, we measured clustering quality using these three complementary measures.

### ARI

The Adjusted Rand Index (ARI)^33^ provides the chance-corrected pairwise agreement between the clustering *C* and the ground truth partition *T*,

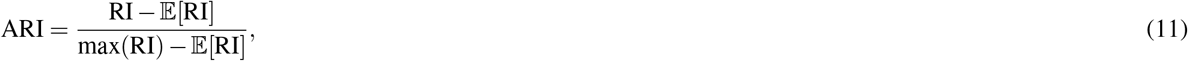

where RI is the Rand index, the fraction of point pairs that are grouped consistently by the two partitions. It ranges from [*−*1, 1], where the worst case score is *−*1, a random clustering scores 0, and the perfect clustering scores 1.

### NMI

The Normalized Mutual Information (NMI)^42^ gives the information shared between the two partitions,

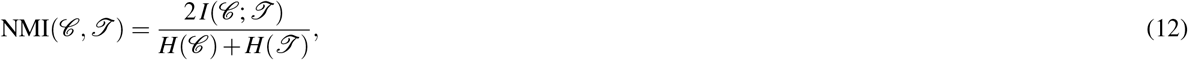

where, *I* is mutual information, and *H* is entropy. It is sensitive to the fragmentation and merging that pairwise measures can miss.

### Hungarian Accuracy

The Hungarian accuracy^43^ gives the identification rate under the best one-to-one mapping between clusters and classes. The mapping is found by the Hungarian algorithm on the co-occurrence matrix **M**, where *M*_*i j*_ counts the images that lie in cluster *i* and class *j*.

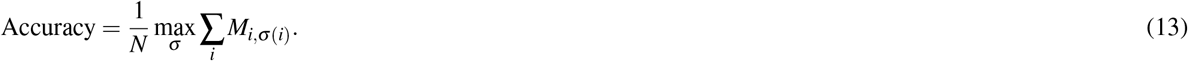

### Annotation Accuracy

Annotation quality was measured as cluster-level and per image accuracy against the folder labels, and the triage was assessed by the per-tier accuracy and by the fraction of images routed to human expert review.

### Declaration of Generative AI and AI-assisted technologies in the writing process

During the preparation of this work, the author(s) used ChatGPT in order to improve grammar and wording. The visualization of the input image grid, dimensionality reduction (PCA and UMAP), and clustering images were generated solely by ChatGPT’s Image 2 model. After using this tool/service, the authors reviewed and edited the content as needed. The author(s) take full responsibility for the content of the publication.

## Results

For model evaluation on the two subsampled datasets, DK-Moth (sub) and HUGO-Bench Aves (sub)^22,41^, every value is the mean over ten random-seed draws and is reported with its standard deviation. EU-Moth and CUB-200-2011 are publicly available datasets, so each was run once. In accordance with the evaluation protocol, noise points identified by density-based methods are excluded from the clustering metrics.

### Species Clustering Across Four Datasets

With the encoder fixed to BioCLIP 2 (ViT-H/14)^6^, we clustered all four datasets using the six algorithms, each at its selected reduction, and report all three metrics per method, along with the clusters found and the noise fraction for the density-based methods (See **Supplementary Material Table S1**).

On the two curated benchmarks, EU-Moth and CUB-200-2011, all six algorithms performed within a narrow range across all metrics. The ARI varied by only 0.031 on EU-Moth (0.955 - 0.986) and 0.019 on CUB-200-2011 (0.867 - 0.886), indicating that the choice of algorithm had little impact on performance. Nevertheless, the density-based methods showed a slight advantage on both datasets. DBSCAN achieved the highest ARI, with 0.986 on EU-Moth and 0.886 on CUB-200-2011, and HDBSCAN achieved the highest Hungarian accuracy on CUB-200-2011 at 0.931. Both methods recovered almost the true number of species and classified less than 1% of EU-Moth and under 3% of CUB-200-2011 images as noise. Overall, on clean imagery, all six methods performed robustly, with only minor differences among them.

The camera-trap subsamples exhibited substantially different behavior, despite both being class-balanced. On DK-Moth (sub), the four fixed-K methods achieved ARI values between 0.684 and 0.715, whereas DBSCAN and HDBSCAN achieved substantially higher ARI values of 0.829 and 0.816, respectively. Thus, the density-based methods outperformed the fixed-K approach by approximately 0.11 ARI, a difference not observed in the curated benchmarks. HUGO-Bench Aves (sub) further accentuated this difference. The fixed-K methods achieved ARI values ranging from 0.394 to 0.588, whereas the density-based methods achieved higher values of 0.830 and 0.853. HDBSCAN, therefore, outperformed the best fixed-K methods by 0.265 ARI (0.853 against 0.588). Since both camera-trap subsamples are class-balanced, class imbalance is unlikely to explain this performance gap. A more plausible explanation is the greater visual variability of camera-trap images, which may be less compatible with the equal-size, hard-partition assumption underlying the fixed-K methods. HUGO-Bench Aves (sub) provides further support for this interpretation: despite containing far fewer species than CUB-200-2011 (30 vs. 200), it yielded lower ARI values for the fixed-K methods. This suggests that image variability and quality, rather than the number of classes, are more important drivers of clustering performance in these datasets. NMI, however, showed smaller differences between methods, as it is less sensitive to disagreements in cluster boundaries. For example, on HUGO-Bench Aves, the advantage of the density-based methods over the best fixed-K method was 0.27 in ARI, compared with 0.12 in NMI (0.906 vs. 0.791).

For both density-based methods, the improved clustering accuracy came at the cost of abstention, with the fraction of images classified as noise increasing with image difficulty. The noise fraction was below 1% on EU-Moth but increased to almost 11% and 18% on the two camera-trap subsamples. Because the evaluation metrics were calculated on the retained images, the density-based methods were not evaluated on quite the same set of samples. DBSCAN consistently discarded more than HDBSCAN, including 16.4% versus 10.8% on DK-Moth (sub). This more aggressive pruning may partly explain much of DBSCAN’s slightly higher ARI, as its performance was evaluated on a smaller and potentially cleaner subset. HDBSCAN retained a larger fraction of the data with only a modest reduction in ARI and achieved the highest ARI on HUGO-Bench Aves at 0.853.

Both density-based methods estimated the number of clusters reasonably well. They found 32 to 34 clusters on HUGO-Bench Aves against a true count of 30, and 233 to 241 on DK-Moth (sub) against 248. Images assigned to low-density regions were therefore left unassigned for potential expert review rather than being forced into an existing cluster. The cluster-selection strategy of HDBSCAN followed the separability of each embedding, with an “excess of mass” on the curated benchmarks and “leaf” on the camera-trap subsamples.

### Foundation Models Comparison

We compared the six foundation models at the pipeline’s intended operation point, clustering the embeddings with HDBSCAN. For a label-free pipeline, HDBSCAN is the natural choice because it infers the number of clusters directly from the data rather than requiring the true number of species as input. It also provides the per-point membership probability, which is used by the uncertainty estimation stage. We ran the comparison on all four datasets. EU-Moth and CUB-200-2011 are single runs, while DK-Moth (sub) and HUGO-Bench Aves (sub) are averaged over ten random subsamples. HDBSCAN labels points that do not belong to sufficiently dense, stable regions as noise. We therefore report the number of recovered clusters and the noise fraction alongside each performance score and interpret these measures jointly. Each backbone is chosen to reduce at a different dimension, which is listed next to it in Table 2.

**Table 2.** Foundation models comparison under HDBSCAN on the four datasets. Each backbone is clustered at its own selected reduction, given as method and target dimension. HDBSCAN uses “excess-of-mass” selection on EU-Moth and CUB-200-2011 and “leaf” on the camera-trap subsamples. DK-Moth (sub) and HUGO-Bench Aves (sub) are mean *±* SD over ten subsamples. EU-Moth and CUB-200-2011 are single fixed runs. ARI, NMI, and Hungarian accuracy are computed on non-noise points, and Noise is the fraction of images HDBSCAN set aside. For the subsampled sets the cluster and noise entries are seed means. Reference partitions contain 200 species for EU-Moth and CUB-200-2011, 248 for DK-Moth (sub), and 30 for HUGO-Bench Aves (sub). **Bold** marks the best backbone in each metric column within a dataset.

| Dataset | Backbone | Reduction | ARI | NMI | Hungarian | Clusters | Noise % |
| --- | --- | --- | --- | --- | --- | --- | --- |
| EU-Moth | BioCLIP 2 (ViT-H/14) | UMAP-20 | **0.983** | **0.995** | **0.992** | 200 | **0.05** |
|  | BioCLIP 2 (ViT-L/14) | UMAP-20 | 0.978 | 0.994 | 0.990 | 200 | 0.09 |
|  | BioCLIP | UMAP-10 | 0.895 | 0.979 | 0.915 | 194 | 2.90 |
|  | DINOv3<sup>†</sup> | UMAP-20 | 0.525 | 0.882 | 0.651 | 181 | 13.92 |
|  | DINOv2<sup>†</sup> | UMAP-20 | 0.458 | 0.843 | 0.577 | 219 | 20.5 |
|  | CLIP | UMAP-15 | 0.425 | 0.827 | 0.559 | 162 | 33.15 |
| DK-Moth (sub) | BioCLIP 2 (ViT-H/14) | UMAP-20 | **0.816 ± 0.018** | **0.955 ± 0.004** | **0.866 ± 0.013** | 241 | **10.8** |
| | BioCLIP 2 (ViT-L/14) | UMAP-20 | 0.727 ± 0.021 | 0.931 ± 0.005 | 0.794 ± 0.013 | 248 | 14.5 |
| | BioCLIP | UMAP-10 | 0.482 ± 0.020 | 0.857 ± 0.006 | 0.616 ± 0.014 | 252 | 20.4 |
| | DINOv3<sup>†</sup> | UMAP-20 | 0.524 ± 0.017 | 0.870 ± 0.006 | 0.648 ± 0.012 | 231 | 25.4 |
| | DINOv2<sup>†</sup> | UMAP-20 | 0.434 ± 0.015 | 0.841 ± 0.004 | 0.578 ± 0.015 | 230 | 26.4 |
| | CLIP | UMAP-15 | 0.273 ± 0.015 | 0.776 ± 0.006 | 0.450 ± 0.012 | 216 | 31.6 |
| CUB-200-2011 | BioCLIP 2 (ViT-H/14) | PCA-150+UMAP-5 | **0.884** | 0.955 | **0.931** | 202 | **1.01** |
|  | BioCLIP 2 (ViT-L/14) | PCA-150+UMAP-5 | 0.882 | **0.956** | 0.926 | 200 | 1.90 |
|  | BioCLIP | PCA-100+UMAP-5 | 0.780 | 0.936 | 0.827 | 190 | 6.17 |
|  | DINOv3<sup>†</sup> | UMAP-5 | 0.716 | 0.920 | 0.765 | 198 | 4.88 |
|  | DINOv2<sup>†</sup> | UMAP-5 | 0.715 | 0.924 | 0.767 | 216 | 4.98 |
|  | CLIP | UMAP-5 | 0.600 | 0.898 | 0.674 | 161 | 17.74 |
| HUGO-Bench Aves (sub) | BioCLIP 2 (ViT-H/14) | UMAP-20 | 0.853 ± 0.033 | 0.906 ± 0.012 | 0.905 ± 0.023 | 32 | 15.4 |
| | BioCLIP 2 (ViT-L/14) | UMAP-20 | 0.851 ± 0.038 | 0.908 ± 0.016 | 0.909 ± 0.020 | 31 | 20.5 |
| | BioCLIP | UMAP-10 | 0.614 ± 0.029 | 0.777 ± 0.010 | 0.719 ± 0.024 | 27 | 34.8 |
|  | DINOv3<sup>†</sup> | UMAP-10 | **0.920 ± 0.013** | **0.942 ± 0.006** | **0.949 ± 0.010** | 31 | **2.6** |
| | DINOv2<sup>†</sup> | UMAP-5 | 0.843 ± 0.015 | 0.906 ± 0.005 | 0.898 ± 0.011 | 32 | 4.7 |
| | CLIP | UMAP-10 | 0.676 ± 0.040 | 0.812 ± 0.014 | 0.751 ± 0.018 | 28 | 34.7 |

<sup>†</sup> DINOv2 and DINOv3 embeddings are taken from the CLS token.

We observed that BioCLIP 2 provided the strongest embedding on three of the four datasets and ranked second on the fourth. Its ViT-H/14 variant achieved the highest ARI on EU-Moth (0.983), DK-Moth (0.816), and CUB-200-2011 (0.884), while DINOv3 performed better on HUGO-Bench Aves (0.920 versus 0.853). The two BioCLIP 2 variants showed similar performance: ViT-L/14 was within 0.005 ARI of ViT-H/14 on EU-Moth, CUB-200-2011, and HUGO-Bench Aves, but 0.089 ARI on the more challenging camera-trap images in the DK-Moth dataset. In contrast, the self-supervised and general-purpose backbones showed substantially less consistent performance across datasets. On the moth datasets, DINOv3 and DINOv2 achieved ARIs of approximately 0.43–0.53, while CLIP performed even worse. On the curated CUB 200-2011 dataset, DINOv3 and DINOv2 achieved ARIs of only 0.716 and 0.715, respectively; whereas on the camera-trap HUGO-Bench Aves dataset, their performance increased to 0.920 and 0.843.

This suggests that DINOv3’s strong performance on HUGO-Bench does not reflect a broadly transferable advantage for bird clustering. On CUB-200-2011, BioCLIP 2 outperformed DINOv3 by 0.168 ARI (0.884 vs. 0.716). Notably, DINOv3’s achieved top performance on HUGO-Bench, the same benchmark on which its authors reported their best results for the model^22^. Thus, our results reproduce the advantage of DINOv3 on HUGO-Bench but indicate that this advantage does not generalize to the other bird dataset evaluated here. The difference in consistency across datasets is particularly pronounced. Across all four datasets, BioCLIP 2 ViT-H/14 achieved ARI scores ranging from 0.816 to 0.983, consistently ranking among the top-performing models. In contrast, DINOv3 ranged from 0.524 to 0.920, while DINOv2 ranged from 0.434 to 0.843.

Thus, the BioCLIP 2 ViT-H/14 model showed more consistent performance across taxa and imaging regimes, suggesting that domain-specific species-level pretraining improves robustness for biological visual tasks. Self-supervised models (DINO) remain strong general-purpose alternatives, but their relative performance is more dataset-dependent. For a tool intended to generalize across arbitrary taxa, this stability is as important as achieving the highest score on any single benchmark. In addition, DINOv2 and DINOv3 do not include text encoders and therefore cannot directly support the subsequent zero-shot annotation stage. BioCLIP 2 (ViT-H/14) was competitive on all four datasets and achieved the highest ARI on three of them. We, therefore, selected BioCLIP 2 ViT-H/14 as the backbone of the proposed pipeline, while DINOv3 remains a strong alternative when clustering performance alone is the primary objective.

The **Supplementary Material Table S2** provides a comparison between our clustering results and those reported by Bjerge et al.^44^ and Markoff et al.^22^ on the EU-Moth, CUB-200-2011, and HUGO-Bench Aves datasets. On these datasets, neither of the two previous approaches achieved higher clustering performance than the method presented in this study.

### Zero-shot Annotation and Confidence-tier Triage

For this step, we kept the BioCLIP 2 and HDBSCAN fixed, achieving cluster-level annotation accuracy of 99.0% on EU-Moth, 97.0% on CUB-200-2011, almost 90% on DK-Moth (sub), and 85.5% on HUGO-Bench Aves (sub). Table 3 shows that zero-shot text matching correctly assigns species labels to nearly all clusters in the two curated benchmarks, identifying 198 of 200 clusters on EU-Moth and 196 of 202 on CUB-200-2011. In contrast, annotation accuracy is lower for the two camera-trap subsampled datasets.

**Table 3.** Cluster-level annotation accuracy and downstream per-image accuracy (non-noise) for the BioCLIP 2 + HDBSCAN pipeline. Camera-trap datasets are mean *±* SD over ten seeds. A cluster counts as correct when its zero-shot label equals its plurality ground-truth species after name normalization.

| Dataset | Images | Clusters | Noise % | Cluster acc. % | Per-image acc. % |
| --- | --- | --- | --- | --- | --- |
| EU-Moth<sup>a</sup> | 2,205 | 200 | 0.05 | 99.0 | 98.23 |
| DK-Moth (sub)<sup>a</sup> | 2,310 | 241 | 10.8 | 89.55 ± 1.5 | 83.91 ± 1.6 |
| CUB-200-2011<sup>b</sup> | 11,788 | 202 | 1.01 | 97.03 | 91.30 |
| HUGO-Bench Aves (sub)<sup>b</sup> | 6,000 | 32 | 15.4 | 85.54 ± 2.14 | 83.21 ± 1.93 |

<sup>a</sup>Labeled with scientific binomials and prompted with the taxonomically structured template (order, family, genus, species) built from GBIF-derived taxonomy.
<sup>b</sup>Labeled with English common names and prompted with the common name.

Per-image accuracy was consistently lower than cluster-level accuracy across all four datasets: 98.2% on EU-Moth, 91.3% on CUB-200-2011, and approximately 83% on both DK-Moth (sub) and HUGO-Bench Aves (sub). This difference can arise from two distinct sources of error: (i) incorrectly named clusters, which propagate an erroneous label to all images assigned to the cluster, and (ii) correctly named clusters that contain images belonging to other species. Table 6 quantifies the contribution of these two error sources.

Table 4 shows that the auto-accept tier achieved high accuracy across all datasets, reaching 98.8% on EU-Moth, 93.2% on CUB-200-2011, 93.1% on DK-Moth (sub), and 88.2% on HUGO-Bench Aves (sub). In contrast, the accuracy in the review tier ranges from 22.8% to 73.0%, while the human-required tier shows even lower accuracy. Thus, across all four datasets, accuracy consistently decreased from the auto-accept tier to the review tier and finally to the human-required tier.

**Table 4.** Confidence-tier accuracy and routing. Each cell gives the within-tier per-image accuracy, the percentage of images in that tier whose predicted species is correct, followed in parentheses by the percentage of the dataset assigned to that tier. The last row is the combined share sent for expert attention (review + human_required). DK-Moth (sub) and HUGO-Bench Aves (sub) are mean *±* SD over ten seeds; EU-Moth and CUB-200-2011 are single runs. Table 5 gives the counts behind every entry, separating each tier into correct, incorrect, and unlabeled images.

| Tier | EU-Moth<br>*n*=2,205 | DK-Moth (sub)<br>*n*=2,310 | CUB-200-2011<br>*n*=11,788 | HUGO-Bench Aves (sub)<br>*n*=6,000 |
| --- | --- | --- | --- | --- |
| auto_accept | 98.8 (98.0) | 93.1 ± 0.8 (75.6 ± 0.9) | 93.2 (95.7) | 88.2 ± 1.2 (78.2 ± 1.2) |
| review | 73.0 (1.7) | 41.7 ± 6.8 (9.5 ± 1.1) | 37.3 (3.2) | 22.8 ± 8.6 (6.4 ± 1.3) |
| human_required | 50.0 (0.3) | 3.7 ± 1.4 (14.9 ± 0.6) | 0.0 (1.2) | 0.0 (15.4 ± 1.9) |
| Expert attention (% of images) | 1.9 | 24.4 | 4.3 | 21.8 |

The distribution of images across the three tiers differed markedly between the curated benchmarks and the camera-trap subsamples. As shown in Figure 3, almost all images in the curated datasets were assigned to auto-accept, with only 1.9% of EU-Moth and 4.3% of CUB-200-2011 routed for expert attention. In contrast, the camera-trap subsamples contained substantially larger fractions of images requiring review, with 24.4% of DK-Moth (sub) and 21.8% of HUGO-Bench Aves (sub) assigned to the review tier.

**Figure 3.**
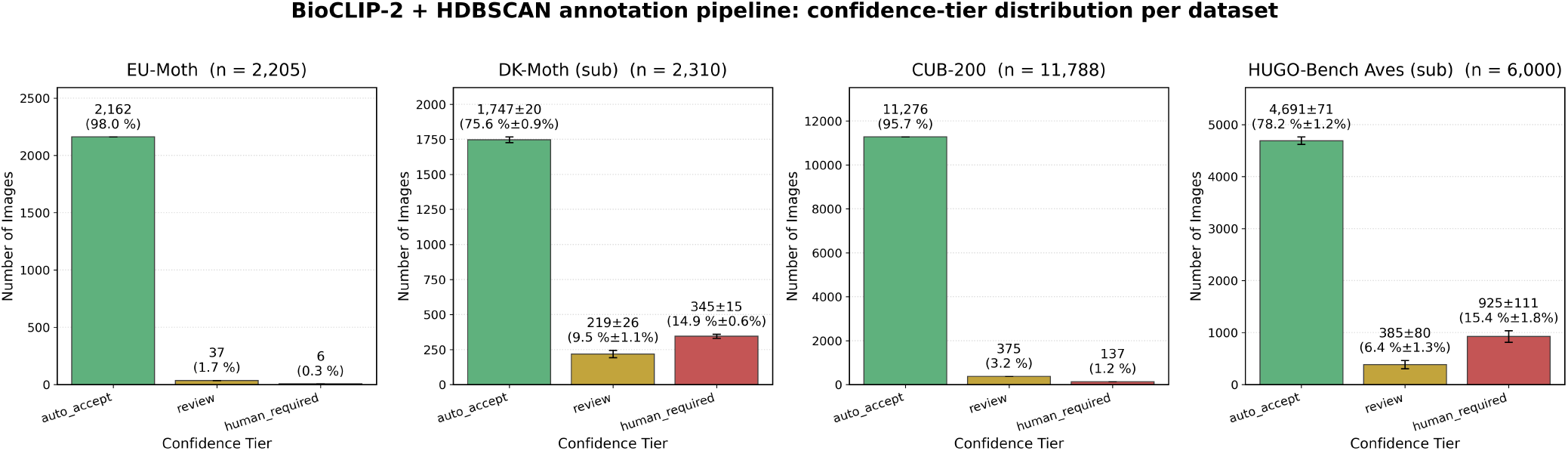
Confidence-tier distribution for the BioCLIP 2 + HDBSCAN pipeline. Number of images routed to each tier per dataset, with the percentage of the dataset in parentheses. DK-Moth (sub) and HUGO-Bench Aves (sub) are mean *±* SD over ten seeds; EU-Moth and CUB-200-2011 are single runs. The two curated benchmarks place almost all images in auto-accept, while the camera-trap subsamples route a larger share to review and human-required. Within-tier accuracy is given in Table 4.

The four-signal score is compared against fixed absolute thresholds rather than a quota, meaning that the proportion of images flagged for review is determined by the difficulty of the collection rather than by a predefined fraction. Images identified as noise by HDBSCAN receive no species prediction, and their uncertainty score is assigned a minimum value of 0.65, regardless of their semantic margin. This exceeds the auto-accept threshold of 0.55, ensuring that noise cannot be auto-accepted. Table 5 shows that across all 22 runs, none of the unclustered images reached the review tier either.

**Table 5.** Composition of each confidence tier, the counts behind Table 4. Every image is named correctly, named incorrectly, or unlabelled, the last being those HDBSCAN left unclustered, which receive no species prediction and so are neither right nor wrong. The three sum to the tier total. EU-Moth and CUB-200 are single runs; DK-Moth (sub) and HUGO-Bench Aves (sub) are mean *±* SD over ten seeds, hence the fractional counts.

|  | EU-Moth<br>*n*=2,205 | DK-Moth (sub)<br>*n*=2,310 | CUB-200<br>*n*=11,788 | HUGO-Bench Aves (sub)<br>*n*=6,000 |
| --- | --- | --- | --- | --- |
| **auto_accept** |  |  |  |  |
| correct | 2,135 | 1,625.8 ± 20.8 | 10,514 | 4,138.2 ± 87.0 |
| incorrect | 27 | 121.0 ± 14.9 | 762 | 552.4 ± 56.3 |
| unlabelled | 0 | 0 | 0 | 0 |
| **review** |  |  |  |  |
| correct | 27 | 90.5 ± 13.8 | 140 | 84.1 ± 29.6 |
| incorrect | 10 | 128.0 ± 25.6 | 235 | 300.7 ± 86.4 |
| unlabelled | 0 | 0 | 0 | 0 |
| **human_required** |  |  |  |  |
| correct | 3 | 12.8 ± 4.6 | 0 | 0 |
| incorrect | 2* | 82.8 ± 14.4 | 18 | 0.2 ± 0.4 |
| unlabelled | 1 | 249.1 ± 20.3 | 119 | 924.4 ± 111.3 |
| Expert attention (% of images) | 1.9 | 24.4 | 4.3 | 21.8 |

\*Neither of these two images shows a moth specimen.

The noises instead accounted for most of the human-required tier on DK-Moth (sub) and CUB-200-2011, and almost all of it on HUGO-Bench Aves (sub). Excluding these images does not improve the reliability of that tier: among the remaining images that carried a species label, only 13.4% were correctly classified on DK-Moth (sub), while none of the 18 labeled images on CUB-200-2011 were correct. Classification errors also persisted within the auto-accept tier, where they were not identified by the uncertainty thresholds. The error rate ranged from 1.2% on EU-Moth to 11.8% on HUGO-Bench Aves. Thus, for the most challenging dataset, more than one in ten auto-accepted images still received an incorrect species label.

Figure 4 shows one image representing each outcome on DK-Moth (sub). The confidently incorrect case illustrates the source of residual error. A cluster of *Dysstroma citrata* was assigned the label *Dysstroma truncata*, although the true species ranked second in the output score rank. The semantic signal, therefore, captured the ambiguity, assigning a score of 0.94 out of 1. However, the other three signals were zero because the cluster was dense, unanimous, and taxonomically consistent. Consequently, the combined score was only 0.33, and all images in the cluster were auto-accepted.

**Figure 4.**
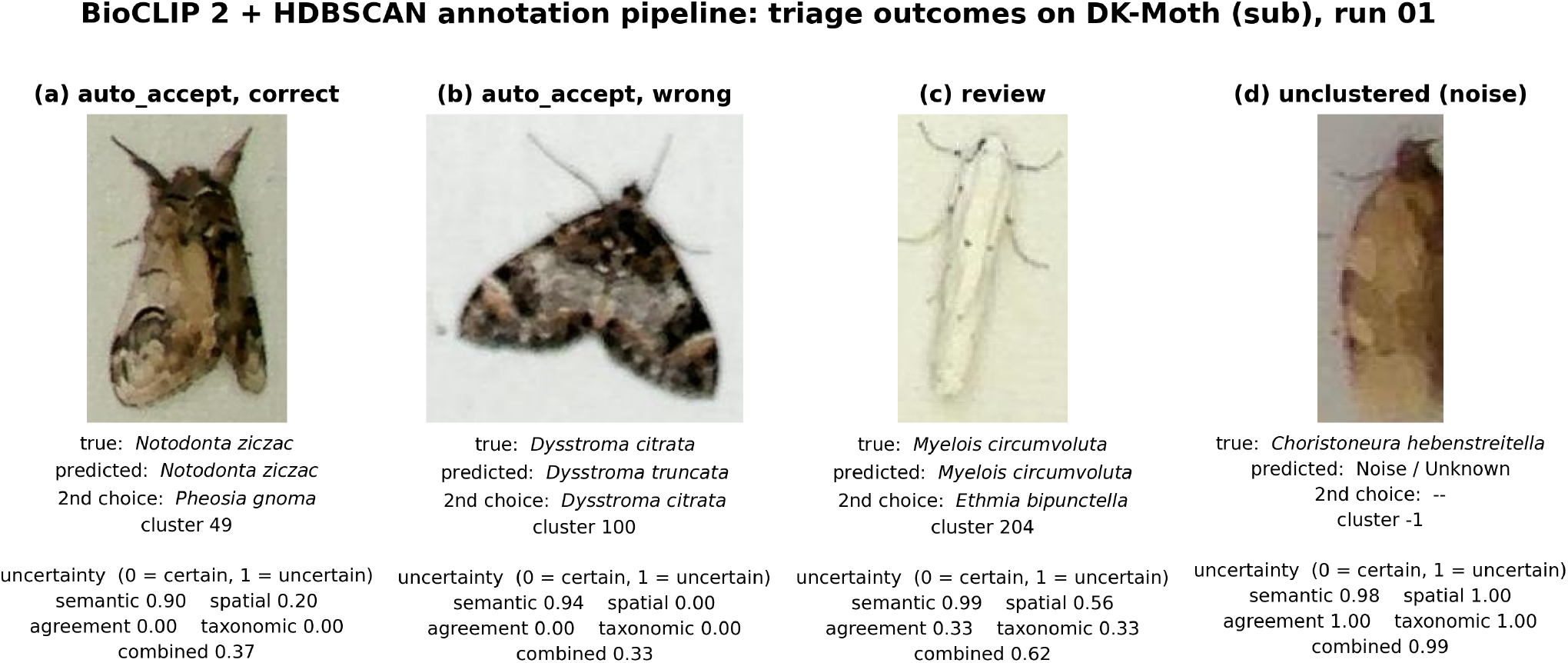
One image from each triage outcome on DK-Moth (sub), run 01 (seed 42). Panels (a), (c), and (d) are the median-scoring image of their category by combined uncertainty, and panel (b) is the lowest-scoring, that is the most confidently wrong, of the images auto-accepted under an incorrect name. In this run the four categories hold 1,648, 118, 212, and 250 images. The four signals are uncertainties on [0, 1], where 0 is certain and 1 is uncertain, and the combined score is their weighted mean with weights 0.35 (semantic), 0.25 (spatial), 0.25 (agreement), and 0.15 (taxonomic). An image is auto-accepted below 0.55, sent to review between 0.55 and 0.70, and to a human at 0.70 or above. In panel (d) the image was never clustered, so it carries no prediction.

This example highlights the structural limitations of the scoring scheme rather than a problem with threshold tuning. The semantic signal has a weight of 0.35, which is below the review threshold of 0.55. Thus, annotation ambiguity alone cannot trigger a review. An image belonging to a tightly formed cluster can, therefore, be auto-accepted even when the assigned species label was selected by only a very narrow semantic margin.

The interactive web report provides partial mitigation. Clusters are organized by tier, with the most urgent cases presented first and ordered within each tier by ascending text margin. Consequently, clusters with the least confident species assignments appear at the top of each section, allowing potentially ambiguous labels to be inspected even when they do not meet the threshold for automatic review.

Auto-accepted errors arise from two distinct sources, which are separated in Table 6. First, the cluster may be assigned an incorrect label, causing every image in the cluster to inherit the error. Second, the cluster may be correctly labeled but contain images belonging to other species, resulting in errors. The first mechanism accounted for most auto-accepted errors on HUGO-Bench Aves (sub), while DK-Moth (sub) showed the opposite pattern, with most errors arising from cluster impurity. Thus, no single mechanism consistently explains the residual error across all four datasets. Because labels are assigned at the cluster level, even a small number of incorrect cluster labels can propagate errors to many images.

**Table 6.**
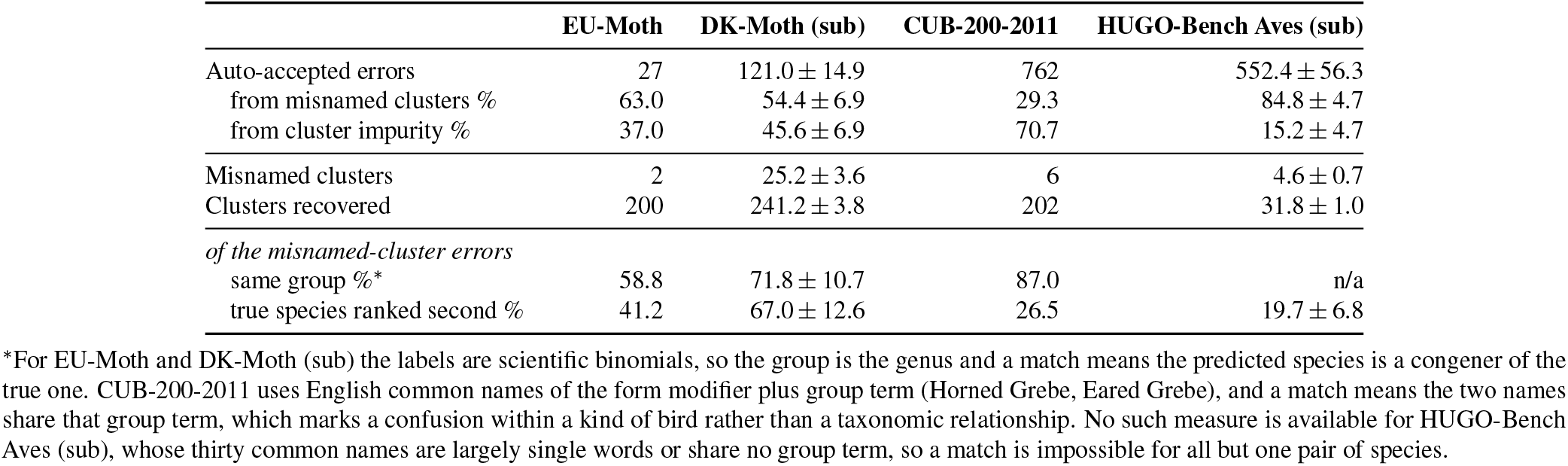
Origin of the auto-accepted errors. An image auto-accepted under a wrong name either sits in a cluster whose label differs from the cluster’s plurality true class, so that every image in it inherits the mistake, or sits in a correctly named cluster and belongs to a different species. The two are mutually exclusive and together cover every wrong auto-accepted image. The last two rows describe only the misnamed-cluster errors, which is the population the naming stage is responsible for. EU-Moth and CUB-200-2011 are single runs; DK-Moth (sub) and HUGO-Bench Aves (sub) are mean *±* SD over ten seeds. The error totals match the auto_accept wrong row of Table 5.

The annotation errors also differed in their nature across datasets. On both moth datasets, the errors were generally near misses, with clusters being assigned the name of a congeneric species. CUB-200-2011 showed a similar pattern at a coarser taxonomic level. Its labels are English common names consisting of a modifier and a group term, and 87% of misnamed clusters were assigned a name from the same group as the true species. Examples include one grebe being labeled as another grebe or one swallow as another swallow. Thus, the model often identified the correct bird group but failed to distinguish between specific species within that group. HUGO-Bench Aves (sub) showed a markedly different pattern. Across all ten runs, none of the misnamed clusters were assigned a name from the same group as the true species. The errors, therefore, represented outright mismatches rather than confusion between visually similar species and can be used to identify clusters of unidentifiable species.

## Discussion

The central question of this study was whether species identification could be automated without requiring the expert effort that the automation is intended to save. This constraint is not merely assumed but has been documented empirically. In a fauna-wide assessment of Hungary, Páll-Gergely et al.^45^ found that almost half of the approximately 36,000 recorded animal species had no active specialist available for identification, with moths identified as one of the groups requiring particularly long training to identify. Supervised classification does not remove this constraint; rather, it shifts the expert workload to an earlier stage, as the labels required for training must still be provided by the same limited pool of specialists.

Our results show that a pipeline composed entirely of pretrained components, without fine-tuning, can correctly name species for between 91.3% and 98.2% of curated images and approximately 83% of camera-trap images. Accuracy alone, however, is not the most important property of the system. The pipeline also provides an indication of which predictions are likely to be reliable and which should be treated with caution, without requiring an expert to review the collection beforehand. We do not claim that these confidence scores are statistically calibrated. Reported confidence and actual correctness are known to diverge in modern neural networks^46^. We therefore report the empirical accuracy achieved at each confidence tier rather than interpreting the confidence scores themselves as calibrated probabilities.

The proportion of a collection that the pipeline can process autonomously depends strongly on image quality. On curated benchmarks, it operates close to full automation, forwarding fewer than 5% of images for further review. In camera-trap collections, it functions primarily as a triage tool, routing approximately one-fifth to one-quarter of images to an expert while still producing incorrect species assignments for approximately 6.9% to 11.8% of auto accepted images. This reduction in performance reflects the greater complexity of field imagery rather than a fundamental limitation of the approach. Camera-trap images exhibit far more variation in pose, illumination, and specimen condition than curated collections, making some performance degradation expected. A method that shows no degradation under these conditions would warrant careful scrutiny rather than confidence. Because the routing is based on absolute confidence thresholds rather than a fixed quota, the size of the review queue itself provides an early indication of collection difficulty before any expert effort is invested. Thus, the practical contribution of the approach is not to eliminate expert involvement but to transform it into a more bounded and predictable workload.

These results have three practical implications for applying the pipeline. The first choice concerns the encoder. DINOv3 produced the best clustering performance on one bird benchmark but dropped to an ARI of 0.524 on moths, whereas BioCLIP 2 achieved ARI values between 0.816 and 0.983 across both taxa. Thus, when the target taxa are not known in advance, a biology-pretrained encoder may provide a more robust choice for both clustering and labeling. The second choice concerns the clustering algorithm. Because the number of species in a survey is generally unknown, and specifying an incorrect number of clusters has little effect on curated data but a substantial effect on camera-trap data, density-based methods may be preferable for uncontrolled imagery. The third choice concerns the appropriate use of the output, which depends on the intended application. For building a verified reference collection, the automatically accepted tier should still be spot-checked, given the error rates reported above. In contrast, for rapid biodiversity surveys or for prioritizing images for expert review, the output can be used directly to support the workflow.

The residual errors are not random, which has important implications for how they can be addressed. In both moth datasets, misnamed clusters typically received the name of a congeneric species, while on CUB-200-2011, 87% were assigned a name from the same common-name group as the true species. Markoff et al. independently observed similar patterns in camera-trap imagery, reporting merged clusters containing “Takahe” with “Australasian Swamphen” (congeneric rails historically treated as conspecific), as well as wolves with black-backed jackals that have similar body proportions^22^. These confusions are consistent with genuine morphological similarity rather than with a defect specific to a particular pipeline.

Two practical consequences follow. First, output that is unreliable at the species level may still be useful for analyzes at the genus or family level, depending on the requirements of the ecological application. Second, when errors are concentrated in cluster naming, correction can be relatively inexpensive because each misnamed cluster represents a single decision rather than multiple individual identifications. This is evident in HUGO-Bench Aves (sub), where fewer than five incorrect cluster names accounted for approximately 85% of the auto-accepted errors. On CUB-200-2011 the errors instead sit inside correctly named clusters, where renaming fixes nothing and each wrong image must be found individually.

It is important to clarify what this work does not resolve. The uncertainty score has a structural blind spot: semantic ambiguity is weighted at 0.35, while review is triggered at 0.55. Consequently, a cohesive and taxonomically consistent cluster assigned the wrong name cannot be flagged on ambiguity alone. The *Dysstroma* case in Figure 4 illustrates the mechanism and accounts for part of the residual error discussed in the Results section. Moreover, all four datasets used in this study have ground-truth labels, meaning that the performance of the confidence tiers was evaluated against known answers. We have not yet tested how the triage system performs when the candidate species list is incomplete or contains incorrect species, as is likely to occur in field applications. Prompting is also restricted to English and relies on an external taxonomy.

The two camera-trap evaluations also used balanced subsamples because, in the full long-tailed DK-Moth collection, the pipeline predominantly recovered well-represented species, while rare classes were either treated as noise or absorbed into larger clusters. Markoff et al. reported a similar failure mode under extreme class imbalance^22^, suggesting that severe class imbalance, rather than collection size itself, is an important condition under which this pipeline degrades.

These limitations point to several directions for further work. Addressing the semantic blind spot will require signals that extend beyond the second-ranked candidate, such as the distribution of similarities across the full candidate list or the similarity between a cluster centroid and the nearest species in the text-embedding space. The combination of signals also warrants further investigation. Because the current uncertainty score is a weighted mean, it allows three confident signals to obscure a fourth that is not, and semantic ambiguity at its current weight can never, on its own, reach the review threshold. Introducing an escalation rule triggered by any single sufficiently uncertain signal could remove this limitation.

Establishing whether the tiers provide reliable guidance in practice will require field evaluation on collections with genuinely unknown compositions, where specialists assess both the species assignments and the routing decisions. At present, expert corrections are used to curate the final dataset but are not fed back into the pipeline. Incorporating these corrections into the signal weights and thresholds could enable an iterative workflow in which subsequent analyzes benefit from decisions already made by the expert^47,48^.

Finally, these directions share an important characteristic of the proposed framework: the encoder is a replaceable component. BioCLIP 2 provides the strongest performance among the models evaluated here, but future foundation models may surpass it. The main contribution of the framework is therefore not a dependence on a particular encoder, but the structure built around it: transforming an initially unlabeled collection into named species groups while explicitly quantifying the remaining need for expert review.

## Data Availability

The DK-Moth dataset is published on Zenodo, and the associated BioCluster code is available on GitHub.

DK-Moth dataset: https://doi.org/10.5281/zenodo.23319153

BioCluster code: https://github.com/sahmedAdnan/biocluster

## Acknowledgements

This research was funded by the Aage V. Jensen Naturfond grant number N2024-0013 and the European Union’s Food, Bioeconomy, Natural Resources, Agriculture and Environment programme, under Grant Agreement ID. 101288886 (BEAGLE). We would like to acknowledge Flemming Helsing-Nielsen, who has labeled the long-tailed DK-Moth camera-trap dataset, and Sikim Chakraborty for his valuable assistance with technical discussions and decision-making throughout the development of this work.

## Author Contributions Statement

S. A.: Conceptualization, Investigation, Methodology, Data curation, Software, Validation, Visualization, Writing – original draft. K. B.: Conceptualization, Investigation, Methodology, Supervision, Writing – review & editing. T. T. H.: Data curation, Funding acquisition, Writing – review & editing. C. N. S. P. : Supervision, Writing – review & editing.

