## Supplementary Material for "BioCluster: Clustering and Zero-Shot Labeling of Ecological Images Using Foundation Models"

This section contains supporting information for comparing the six clustering algorithms and related work performing clustering on the datasets EU-Moth, CUB-200-2011, and HUGO-Bench Aves.

### Species Clustering Across Four Datasets

With the encoder fixed to BioCLIP 2 (ViT-H/14)[1], we clustered all four datasets using the six clustering algorithms, each at its selected reduction, and report all three metrics per method, along with the clusters found and the noise fraction for the density-based methods; see Table S1, DBSCAN, and HDBSCAN.

**Table. S1.** Per-method clustering metrics for BioCLIP 2 (ViT-H/14) on the four datasets. The reduction (mode and target dimension) selected for each dataset is shown beneath its name. Clusters and noise apply to the density-based methods. ARI, NMI, and Hungarian accuracy are computed without noise points. The fixed-K methods use the known species count (shown) and produce no noise (–). DK-Moths (sub) and HUGO-Bench Aves (sub) metrics are mean ± SD over ten seeds, and their cluster and noise entries are seed means. **Bold** marks the best score within each dataset.

| Dataset | Method | ARI | NMI | Hungarian | Clusters | Noise % |
|---|---|---|---|---|---|---|
| EU-Moth | | | | | | |
| UMAP-20D | K-means | 0.980 | 0.994 | 0.990 | 200 | – |
| | Agglomerative | 0.979 | 0.994 | 0.990 | 200 | – |
| | Spectral | 0.955 | 0.989 | 0.973 | 200 | – |
| | GMM | 0.980 | 0.994 | 0.990 | 200 | – |
| | DBSCAN | **0.986** | **0.996** | **0.993** | 200 | 0.91 |
| | HDBSCAN (eom) | 0.983 | 0.995 | 0.992 | 200 | 0.05 |
| DK-Moth (sub) | | | | | | |
| UMAP-20D | K-means | 0.714 ± 0.013 | 0.926 ± 0.003 | 0.787 ± 0.012 | 248 | – |
| | Agglomerative | 0.715 ± 0.010 | 0.928 ± 0.002 | 0.785 ± 0.008 | 248 | – |
| | Spectral | 0.684 ± 0.013 | 0.916 ± 0.003 | 0.758 ± 0.012 | 248 | – |
| | GMM | 0.712 ± 0.015 | 0.926 ± 0.004 | 0.785 ± 0.013 | 248 | – |
| | DBSCAN | **0.829 ± 0.012** | **0.965 ± 0.002** | **0.867 ± 0.007** | 233 | 16.44 |
| | HDBSCAN (leaf) | 0.816 ± 0.018 | 0.955 ± 0.004 | 0.866 ± 0.013 | 241 | 10.78 |
| CUB-200-2011 | | | | | | |
| PCA-150D+UMAP-5D | K-means | 0.867 | 0.953 | 0.909 | 200 | – |
| | Agglomerative | 0.867 | 0.953 | 0.908 | 200 | – |
| | Spectral | 0.872 | 0.952 | 0.925 | 200 | – |
| | GMM | 0.868 | 0.953 | 0.910 | 200 | – |
| | DBSCAN | **0.886** | **0.957** | 0.916 | 226 | 2.84 |
| | HDBSCAN (eom) | 0.884 | 0.955 | **0.931** | 202 | 1.01 |
| HUGO-Bench Aves (sub) | | | | | | |
| UMAP-20D | K-means | 0.588 ± 0.006 | 0.791 ± 0.008 | 0.760 ± 0.012 | 30 | – |
| | Agglomerative | 0.584 ± 0.019 | 0.787 ± 0.009 | 0.750 ± 0.014 | 30 | – |
| | Spectral | 0.394 ± 0.046 | 0.730 ± 0.028 | 0.609 ± 0.075 | 30 | – |
| | GMM | 0.575 ± 0.014 | 0.780 ± 0.005 | 0.733 ± 0.021 | 30 | – |
| | DBSCAN | 0.830 ± 0.018 | 0.890 ± 0.005 | 0.880 ± 0.009 | 34 | 17.35 |
| | HDBSCAN (leaf) | **0.853 ± 0.033** | **0.906 ± 0.012** | **0.905 ± 0.023** | 32 | 15.40 |

### Comparison with Existing Clustering Methods

Two recent pipelines are sufficiently similar to ours to permit a meaningful comparison. Bjerge et al.[2] fine-tuned ImageNet-pretrained backbones using episodic meta-learning and subsequently clustered the resulting feature representations. They reported results on EU-Moth and CUB-200-2011, the two benchmarks we evaluated in this study, as shown in Table S2. However, the evaluation protocols differ in two important respects. First, their few-shot setting clustered only the held-out test classes (20 on EU-Moth and 40 on CUB-200-2011), whereas we cluster all 200 species in each benchmark. Second, their backbone was fine-tuned for the task, whereas we use an unmodified encoder strictly as a fixed feature extractor.

On EU-Moth, our method achieved an ARI of 0.983 compared to 0.897 for Bjerge et al.[2], while the methods performed nearly identically on CUB-200-2011 (0.884 vs. 0.881). NMI showed the same overall pattern (0.995 vs. 0.962) on EU-Moth and 0.955 vs. 0.932 on CUB-200-2011. However, NMI is less informative for this comparison because it is not adjusted for chance and generally increases with the number of clusters; our partitions contain substantially more clusters because they resolve all species rather than only the held-out subset. The ARI comparison is therefore more appropriate. By this measure, BioCLIP 2 (ViT-H/14) matches or exceeds the performance of the fine-tuned approach while clustering five to ten times as many species.

**Table. S2. Clustering quality against the closest fine-tuned method on the two shared benchmarks**, on the two metrics both systems report. Our entries are HDBSCAN at the reduction selected for each dataset. The reference entries are the best configuration reported by Bjerge et al.[2]. **Bold** marks the higher value per metric within each benchmark.

| Benchmark | System | Classes | Fine-tune | NMI | ARI |
|---|---|---|---|---|---|
| EU-Moth | ConvNeXt-B[2] | 20 | Episodic | 0.962 | 0.897 |
| EU-Moth | This work (BioCLIP 2) | 200 | None | **0.995** | **0.983** |
| CUB-200-2011 | ConvNeXt-B[2] | 40 | Classic | 0.932 | 0.881 |
| CUB-200-2011 | This work (BioCLIP 2) | 200 | None | **0.955** | **0.884** |

Markoff et al.[3] also evaluated several backbone models and benchmarked zero-shot clustering, in their case, using the HUGO-Bench camera-trap dataset. We evaluate the same 30-species Aves partition under the same protocol of 200 images per species. Thus, the two sets of results are based on identical images and directly comparable evaluation protocols. The evaluation metrics are also identical. V-measure is the normalized mutual information taken with the arithmetic mean of the two entropies, which is what our NMI column reports. Their strongest unsupervised result on the birds is DINOv3 at 0.948, reached by projecting to two dimensions with t-SNE and clustering with HDBSCAN, recovering 33 clusters and setting aside 0.9% of the images as outliers. Our DINOv3 reaches $0.942 \pm 0.006$ on the same birds with 31 clusters and 2.6% noise, and is likewise the strongest of the six embeddings we tested. The two studies agree on the ranking.

The two studies differ in their findings for BioCLIP 2. In their benchmark, BioCLIP 2 scored 0.730, placing it between DINOv2 and CLIP, whereas on the bird partition in our setup it matches DINOv2, with scores of $0.908 \pm$,0.016 versus $0.906 \pm$,0.005, and DINOv3 outperforms both. The pipelines differ in their dimensionality reduction and clustering parameters, and their evaluation pools birds together with mammals, so we do not attribute the discrepancy to any single design choice. What both studies agree on is the overall ranking pattern rather than the exact gaps between models. DINOv3 leads on this partition, and CLIP ranks near the bottom in both cases, while the relative ordering of the intermediate models shifts with the configuration. This suggests that an encoder should be selected jointly with its reduction and clustering settings rather than based solely on a published ranking. The more substantial difference lies in scope. Their pipeline ends at the clustering stage and forwards its outliers for expert review. In contrast, our pipeline proceeds further by assigning a name to each cluster and grading every image using the four-signal triage. We revisit the benefits of these additional stages in the Discussion.
